# The *yellow* gene influences *Drosophila* male mating success through sex comb melanization

**DOI:** 10.1101/673756

**Authors:** Jonathan H. Massey, Daayun Chung, Igor Siwanowicz, David L. Stern, Patricia J. Wittkopp

## Abstract

*Drosophila melanogaster* males perform a series of courtship behaviors that, when successful, result in copulation with a female. For over a century, mutations in the *yellow* gene, named for its effects on pigmentation, have been known to reduce male mating success. Prior work has suggested that *yellow* influences mating behavior through effects on wing extension, song, and/or courtship vigor. Here, we rule out these explanations, as well as effects on the nervous system more generally, and find instead that the effects of *yellow* on male mating success are mediated by its effects on pigmentation of male-specific leg structures called sex combs. Loss of *yellow* expression in these modified bristles reduces their melanization, which changes their structure and causes difficulty grasping females prior to copulation. These data illustrate why the mechanical properties of anatomy, and not just neural circuitry, must be considered to fully understand the development and evolution of behavior.

## Introduction

> “*The form of any behavior depends to a degree on the form of the morphology performing it*”
>
> — -Mary Jane West-Eberhard, 2003

Over 100 years ago in Thomas Hunt Morgan’s fly room, Alfred Sturtevant described what is often regarded as the first example of a single gene mutation affecting behavior (Sturtevant, 1915; reviewed in Drapeau *et al*., 2003; Cobb, 2007; Greenspan 2008): he noted that *yellow* mutant males, named for their loss of black pigment that gives their body a more yellow appearance (Figure 1A), mated successfully with wild-type females much less often than wild-type males. In 1956, in what is often regarded as the first ethological study (reviewed in Cobb, 2007; Greenspan 2008), Margaret Bastock compared courtship of *yellow* mutant and wild-type males and concluded that despite all courtship actions being present, loss of *yellow* function likely reduces courtship vigor or drive, leading to copulation inhibition (Bastock 1956). Despite more recent data consistent with this hypothesis (Drapeau et al. 2003), the precise mechanism by which the *yellow* gene affects male mating success in *D. melanogaster* has remained a mystery. Consequently, Bastock’s statement about *yellow* from her 1956 paper is equally true today: *“It seemed worthwhile therefore to examine more closely one example of a gene mutation affecting behavior and to ask two questions, (1) how does it bring about its effect? [and], (2) what part might it play in evolution?”*

**Fig. 1.**
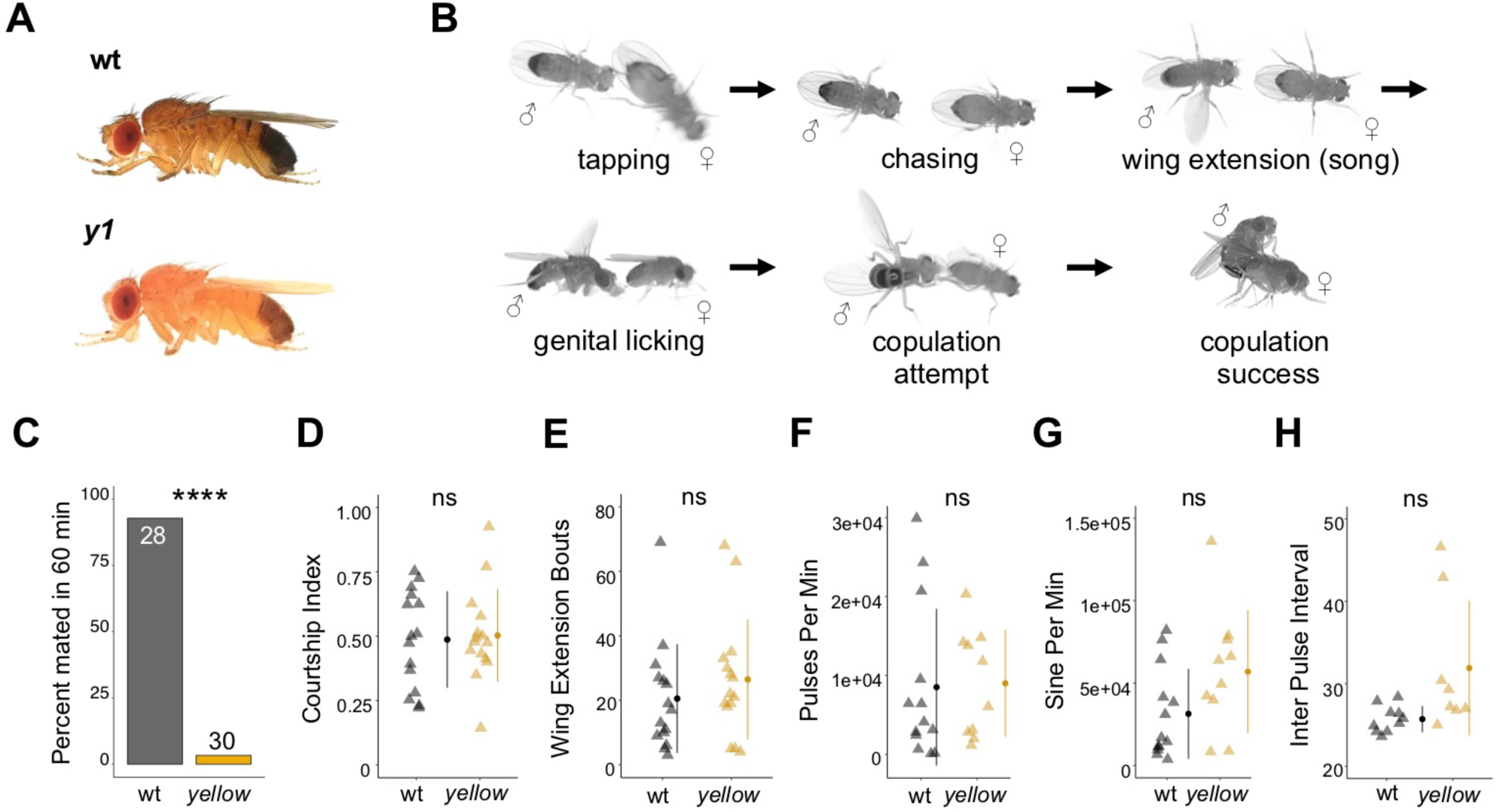
The *Drosophila melanogaster yellow* gene is required for male mating success. (**A**) Photographs comparing wild-type and *yellow* (*y1*) body pigmentation (Nicolas Gompel). (**B**) Snapshots from videos illustrating *D. melanogaster* courtship behaviors. (**C**) *y1* males (yellow) showed significantly lower mating success levels compared to wild-type males (black) in non-competitive, one-hour trials. Sample sizes are shown at the top of each barplot. (**D**-**H**) *y1* males showed similar levels of courtship activity and song compared to wild-type males. (**D**) Courtship index: the proportion of time a male engages in courtship activity divided by the total observation period. (**E**) Wing extension bouts: the number of unilateral wing extensions during the observation period. (**F**) Pulses per minute. (**G**) Sine per minute. (**H**) Inter pulse interval. (**D-H**) Show individual points that represent single fly replicates. Circles represent means and lines SD. Significance was measured using Fisher’s exact test in (**C**), Student’s t-tests (two-tailed) in (**D**,**E**), and one-way ANOVA in (**F**-**H**). \*\*\*\**P*<0.0001. n.s., not significant.

The *D. melanogaster yellow* gene encodes a protein hypothesized to act either structurally (Geyer *et al*., 1986) or enzymatically (Wittkopp *et al*., 2002) in the synthesis of dopamine melanin, and a Yellow homolog has been shown to bind dopamine and other biogenic amines in the sand fly *Lutzomyia longipalpis* (Xu *et al*., 2011). The interaction between Yellow and dopamine might explain the protein’s effects on male mating success because dopamine acts as a modulator of male courtship drive in *D. melanogaster* (Zhang *et al*., 2016). These effects of dopamine are mediated by neurons expressing the gene *fruitless* (*fru*) (Zhang *et al*., 2016), which is a master regulator of sexually dimorphic behavior in *D. melanogaster* that can affect every component of courtship and copulation (reviewed in Villella and Hall, 2008). *fru* has also been shown to regulate expression of *yellow* in the central nervous system (CNS) of male *D. melanogaster* larvae (Drapeau *et al*., 2003). These observations suggest that the pleiotropic effects of *yellow* on male mating success might result from effects of *yellow* in the adult CNS, particularly in *fru*-expressing neurons. Consistent with this hypothesis, functional links between the pigment synthesis pathway and behavior mediated by the nervous system have previously been reported for other pigmentation genes (Hotta and Benzer, 1969; Heisenberg, 1971; Borycz *et al*., 2002; Richardt *et al*., 2002; True *et al*., 2005; Suh and Jackson, 2007).

## Results and Discussion

### fruitless*-expressing cells do not mediate the effect of* yellow *on male mating success*

*D. melanogaster* males perform multiple behaviors, including tapping, chasing, singing, and genital licking, before attempting to copulate with females by curling their abdomen and grasping the female (Figure 1B, Movie 1). In one-hour trials, we found that virgin males homozygous for a null allele of the *yellow* gene (*y1*) successfully mated with wild-type virgin females only 3% of the time, whereas wild-type males mated with wild-type virgin females 93% of the time (Figure 1C). Videos of mating trials indicated that the difference in mating success between wild-type and *yellow* males did not come from differences in courtship activity (Figure 1D-H) (compare Movies 1 and 2), but rather from differences in the ability of *yellow* and wild-type males to initiate copulation (compare Movies 3 and 4).

To determine whether *yellow* activity in *fru*-expressing cells is responsible for this difference in mating success, we used the UAS-GAL4 system (Brand and Perrimon, 1993) to drive expression of *yellow*-*RNAi* (Dietzl *et al.*, 2007) with *fru*^*GAL4*^ (Stockinger *et al*., 2005), knocking down native *yellow* expression in these cells. We also used *fru*^*GAL4*^ to drive *yellow* expression in *y1* mutants. In both cases, we found no significant effect on male mating success (Figure 2A,B), showing that expression of *yellow* in *fru*-expressing cells is neither necessary nor sufficient for *yellow’s* effect on male mating success.

**Fig. 2.**
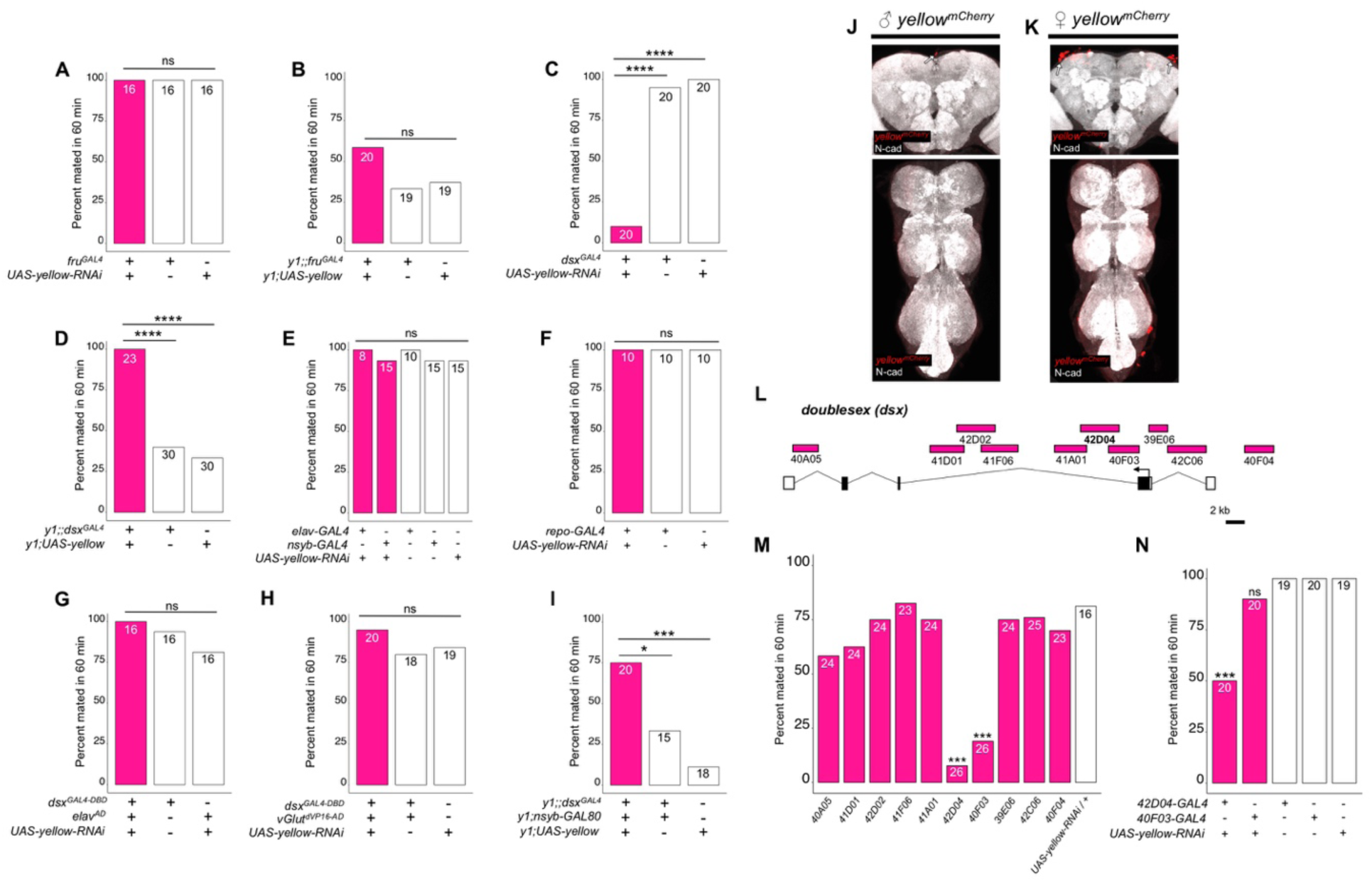
*yellow* expression in non-neuronal *doublesex*-expressing cells, but not *fruitless*-expressing cells, is necessary and sufficient for male mating success. **(A**,**B**) Neither expressing *yellow-RNAi* nor *yellow-cDNA* in *fru*-expressing cells using *fru*^*GAL4*^ (Stockinger *et al*., 2005) affected male copulation. (**C**) Expressing *yellow-RNAi* in *dsx*-expressing cells using *dsx*^*GAL4*^ (Robinett *et al*., 2010) significantly inhibited male mating success. (**D**) Expressing *yellow* in *dsx*-expressing cells using *dsx*^*GAL4*^ in a *y1* mutant background was sufficient to restore male mating success. (**E**,**F**) Expressing *yellow-RNAi* using pan-neuronal (*elav-GAL4* and *nsyb-GAL4*) and pan-glia (*repo-GAL4*) drivers did not affect male mating success. (**G**) Restricting *yellow-RNAi* expression to *dsx*-expressing neurons using the split-GAL4 technique, combining *dsx*^*GAL4-DBD*^ (Pavlou *et al*., 2016) with *elav*^*VP16-AD*^ (Luan *et al*., 2006), did not affect male mating success. (**H**) Restricting *yellow-RNAi* expression to *dsx*-expressing glutamatergic neurons using the split-GAL4 technique, combining *dsx*^*GAL4-DBD*^ (Pavlou *et al*., 2016) with *vGlut*^*dVP16-AD*^ (Gao *et al*., 2008) did not affect male mating success. (**I**) Expressing *yellow* in *dsx*-expressing cells restricted outside the CNS using *dsx*^*GAL4*^ and *nsyb-GAL80* (courtesy of Julie Simpson) in a *y1* mutant background significantly increased male mating success. (**J**,**K**) Brain and ventral nerve cord of adult male and female *y*^*mCherry*^ flies stained with anti-N-Cadherin (N-cad) antibody labeling neuropil (white) and anti-DsRed antibody labeling Yellow::mCherry (red). We observed sparse, inconsistent signal outside the CNS at the top of the brain in males (white arrow), and especially females (white arrow), but we were unable to confirm a previous report that *y*^*mCherry*^ is expressed in the adult brain (Hinaux *et al*., 2018). (**L**) Diagram of the male exon structure of the *dsx* locus highlighting 10 genomic fragments between 1.7 and 4 kb used to clone Janelia enhancer trap GAL4 drivers (Pfeiffer *et al*., 2008). Black boxes indicate coding exons. White boxes indicate 5’ and 3’ UTRs, and the arrow in exon 2 denotes the transcription start site. (**M**) Expressing *yellow-RNAi* using each Janelia *dsx-GAL4* driver identified *42D04-GAL4* and *40F03-GAL4* as affecting male mating success when compared with the *yellow-RNAi* control. (**N**) A replicate experiment comparing *42D04-GAL4* and *40F03-GAL4* effects on male mating success with both GAL4 and UAS parental controls confirmed the significant effect of *42D04-GAL4* but not *40F03-GAL4*. We attribute differences in the *40F03-GAL4* effect between (**M**) and (**N**) to between experiment variability in the levels of male mating success; each common genotype tested in (**M**), for example, mated at higher levels in (**N**), but *42D04-GAL4* consistently showed a significant effect relative to controls. Sample sizes are shown at the top of each barplot. Significance was measured using Chi-square tests with Bonferroni corrections for multiple comparisons. \**P*<0.05, \*\*\**P*<0.00, \*\*\*\**P*<0.0001. n.s., not significant.

### Doublesex-expressing cells require *yellow* for normal male mating success

To continue searching for cells responsible for *yellow*’s effects on mating, we examined a 209 bp sequence 5’ of the *yellow* gene called the “mating-success regulatory sequence” (MRS) because deletion mapping indicated it was required for male mating success (Drapeau et al. 2006). We hypothesized that the MRS might contain an enhancer driving *yellow* expression and found that ChIP-seq data indicates the Doublesex (Dsx) transcription factor binds to this region *in vivo* (Clough *et al*., 2014). Like *fru, dsx* expression is required to specify sex-specific behaviors in *D. melanogaster* (Rideout *et al*., 2010; Robinett *et al*., 2010; reviewed in Villella and Hall, 2008; Yamamoto and Koganezawa, 2013), suggesting that *yellow* expression regulated by Dsx through the MRS enhancer might be responsible for its effects on male mating behavior. We found that reducing *yellow* expression in *dsx*-expressing cells with either of two different *dsx*^*GAL4*^ drivers (Robinett *et al*., 2010; Rideout *et al*., 2010) strongly reduced male mating success (Figure 2C, Supplementary Figure S1A), whereas restoring *yellow* activity in cells expressing *dsx*^*GAL4*^ in *y*^*1*^ mutants significantly increased male mating success compared with *y*^*1*^ controls (Figure 2D, Supplementary Figure S1B). Video recordings of male flies with reduced *yellow* expression in *dsx*-expressing cells showed the same mating defect observed in *y*^*1*^ mutants: males seem to perform all courtship actions normally, but repeatedly failed to copulate (Movie 5). We therefore conclude that *yellow* expression is required in *dsx*-expressing cells for normal male mating behavior.

To determine whether the MRS sequence might be the enhancer mediating *yellow* expression in *dsx*-expressing cells that affect male mating success, we manipulated *yellow* expression with GAL4 driven by a 2.7kb DNA region located 5’ of *yellow* that includes the wing, body, and putative MRS enhancers (Gilbert *et al*., 2006, Supplementary Figure S2A). Altering *yellow* expression with this GAL4 driver modified pigmentation as expected but did not affect male mating success (Supplemental Figure S2B-D), possibly because this GAL4 line did not show any detectable expression in the adult CNS (Supplementary Figure S2E). To test more directly whether the MRS was necessary for male mating success, we deleted 152 bp of the 209 bp MRS sequence using CRISPR/Cas9 gene editing (Bassett *et al*., 2013) (Supplemental Figure S2F,G). We found that this deletion had no significant effect on male mating success (Supplemental Figure S2H), contradicting the previous deletion mapping data (Drapeau *et al*., 2006). We conclude therefore that *yellow* expression in *dsx*-expressing cells affecting mating behavior must be mediated by other *cis*-regulatory sequences associated with the *yellow* gene.

### dsx*-expressing cells outside the CNS require* yellow *for normal male mating success*

Although *dsx* is expressed broadly throughout the fly (Robinett *et al*., 2010; Rideout *et al*., 2010), we hypothesized that its expression in the nervous system would be responsible for *yellow*’s effects on mating because *yellow* has been reported to be expressed in the adult brain (Hinaux *et al*., 2018) and behavioral effects of other pigmentation genes are mediated by neurons (Hotta and Benzer, 1969; Heisenberg, 1971; Borycz et al., 2002; True *et al*., 2005). However, we found that suppressing *yellow* expression in the larval CNS, dopaminergic neurons, or serotonergic neurons (Supplementary Figure S3), or in all neurons (Figure 2E) or all glia (Figure 2F), had no significant effect on male mating success. Specifically reducing *yellow* expression in either all *dsx*-expressing neurons (Figure 2G) or all *dsx*-expressing glutamatergic neurons that are required for genital coupling (Pavlou *et al*., 2016) (Figure 2H) also had no significant effect on male mating success. In addition, when we examined *yellow* expression in adult brains, we were only able to observe non-specific signal at the anterior of the adult brain in females (Figure 2J,K). Given this lack of evidence that *yellow* is required in neuronal cells for normal male mating behavior, we limited *dsx*^*GAL4*^ activation of *yellow* expression in *y1* mutants to non-neuronal cells and found that these flies exhibited a substantial increase in male mating success compared with *y*^*1*^ mutant males (Figure 2I), showing that *yellow* expression in non-neuronal *dsx*-expressing cells is required for normal male mating behavior.

To identify which non-neuronal *dsx*-expressing cells require *yellow* expression for normal male mating success, we screened ten *dsx*-enhancer GAL4 lines that each contains a different ~3 kb region of *dsx* noncoding sequence (Figure 2L; Pfeiffer *et al*., 2008). Two of these lines, *42D04-GAL4* and *40F03-GAL4*, significantly decreased male mating success when driving *yellow*-*RNAi* (Figure 2M). These two GAL4 drivers contain overlapping sequences from intron 2 of *dsx* (Figure 2L), suggesting that their similar effects result from reduction of *yellow* expression in the same cells. Line *42D04-GAL4* had stronger effects than *40F03-GAL4* (Figure 2N), so we performed all further analyses with this line. Males with *yellow* reduced by *42D04-GAL4* performed courtship behavior in a pattern similar to *y*^1^ mutant males: males performed all precopulatory courtship behaviors normally, but repeatedly failed to copulate, even after hours of attempts (Movie 6). These data indicate that some or all cells in which *42D04-GAL4* drives expression require *yellow* expression for normal male mating behavior.

### Sex combs require *yellow* expression for normal male mating success

*42D04-GAL4* drives expression in a sexually dimorphic pattern in multiple neurons of the adult male (Figure 3A,B) and female CNS (Supplemental Figure S4A,B), consistent with previously described *dsx*^*GAL4*^ expression in the posterior cluster, the abdominal cluster, and, in males, in the prothoracic TN1 neurons (Robinett *et al*., 2010). *42D04-GAL4* also drives expression in male and female larval CNS and genital discs, with expression in the genital tissues persisting into the adult stage only in females (Supplemental Figure S4C-G). Finally, we observed *42D04-GAL4* expression at the base of the sex combs (also observed by Robinett *et al*. 2010), which are modified bristles used during mating (Cook, 1975; Ng and Kopp 2008; Hurtado-Gonzales *et al*., 2015) that are present only on the first tarsal segment of adult male forelegs (Figure 3C-F). Yellow protein is expressed in sex combs (Hinaux *et al*., 2018, Figure 3G,H), where it is presumably required for synthesis of black dopamine melanin in the sex comb “teeth”. This expression of *yellow* in sex comb cells is driven by enhancer sequences in the *yellow* intron (Supplementary Figure S5), potentially explaining why manipulating *yellow* expression using GAL4 driven by sequences 5’ of the *yellow* gene failed to affect mating. Driving expression of *yellow*-RNAi with *42D04-GAL4* eliminated expression of an mCherry tagged version of the native Yellow protein in sex combs and strongly reduced black melanin in the sex combs (Figure 3I-L) but not the abdomen (Supplemental Figure S4J).

**Fig. 3.**
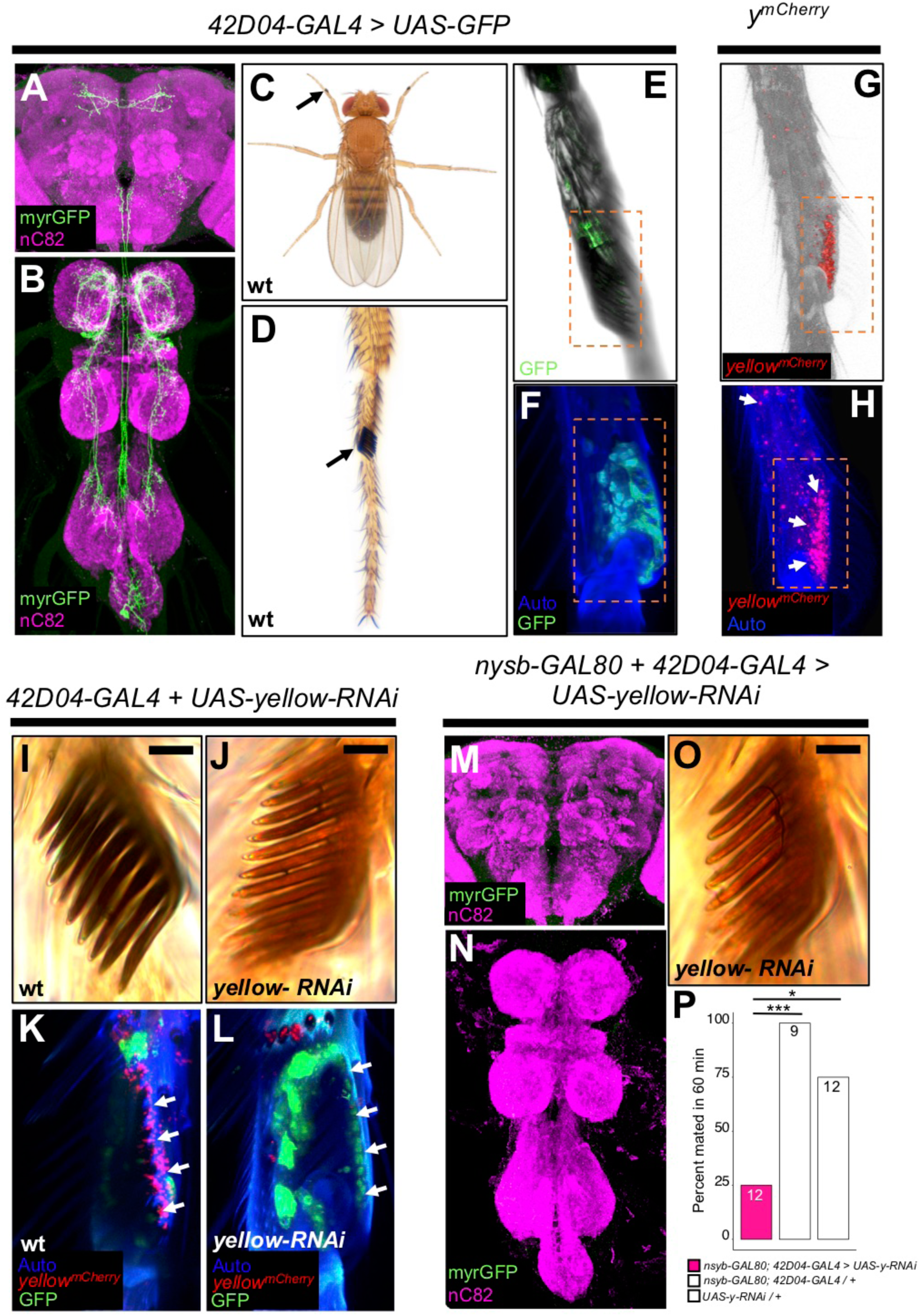
*yellow* expression in non-neuronal *42D04-GAL4* expressing cells is necessary for sex comb melanization and male mating success. (**A**,**B**) Brain and ventral nerve cord of adult male fly stained with anti-GFP (green) antibody for myrGFP expressed using *42D04-GAL4* and counterstained with anti-nC82 (magenta) for neuropil. (**C**) Wild-type (wt) *D. melanogaster* adult male fly highlighting the location of sex combs (Nicolas Gompel). (**D**) Close up of a wild-type (wt) sex comb on the first tarsal segment (ts1) of the front leg (courtesy of Nicolas Gompel). (**E**) Bright field illumination of a male front leg expressing cytGFP (green) in sex-comb cells using *42D04-GAL4*. (**F**) Confocal image of the sex comb cells expressing cytGFP (green) with *42D04-GAL4* and leg cuticle autofluorescence (blue). (**G**) Confocal image of a *y*^*mCherry*^ male leg highlighting native *y*^*mCherry*^ sex comb expression (red). (**H**) Zoomed in confocal image shown in (**G**) with leg cuticle autofluorescence (blue) and native *y*^*mCherry*^ sex comb expression (red). (**I**) Wild-type (wt) sex comb. (**J**) Loss of black melanin in sex combs in males expressing *yellow-RNAi* using *42D04-GAL4.* (**K**) Co-localization of *y*^*mCherry*^ (red) at the base of the sex comb cells expressing cytGFP (green) with *42D04-GAL4*. (**L**) Loss of *y*^*mCherry*^ (red) at the base of the sex comb cells expressing cytGFP (green) and *yellow-RNAi* using *42D04-GAL4*. (**M**,**N**) Brain and ventral nerve cord of adult male expressing *nsyb-GAL80* to block GAL4 activity in the CNS, stained with anti-GFP (green) antibody for myrGFP expressed using *42D04-GAL4*, and counterstained with anti-nC82 (magenta) for neuropil. (**O**) Loss of black melanin in sex combs in *nsyb-GAL80* males expressing *yellow-RNAi* using *42D04-GAL4*. (**P**) Expressing *yellow-RNAi* using *42D04-GAL4* in males expressing *nsyb-GAL80* significantly inhibited male mating success. Scale bars in (**I**), (**J**), and (**O**) measure 12.5 μm. Sample sizes are shown at the top of each barplot. Significance was measured using Chi-square tests with Bonferroni corrections for multiple comparisons. \**P*<0.05, \*\*\**P*<0.001.

To test the impact of *yellow* expression in sex combs on male mating behavior, we used *42D04-GAL4* to drive *yellow-RNAi*, but inhibited the function of *42D04-GAL4* in the CNS with *nysb-GAL80* (courtesy of Julie Simpson). These flies showed no GAL4 activity in the CNS (Figure 3M,N), but lost black melanin in the sex combs (Figure 3O) and had significantly reduced male mating success (Figure 3P). High-speed videos (1000 frames per second) revealed that *yellow* mutant (*y*^1^) males fail repeatedly to grasp the female abdomen with their sex combs when attempting to mount and copulate (Movie 7), whereas wild-type males more readily grasp the female with their melanized sex combs and initiate copulation efficiently (Movie 8). These observations suggest that *yellow* expression in sex combs affects their melanization, which in turn affects their function.

### Sex comb melanization is required for efficient grasping, mounting and copulation

To test whether sex comb melanization (as opposed to some other unknown effect of losing *yellow* expression in sex combs) is critical for male sexual behavior, we suppressed expression of *Laccase2* (Arakane *et al*., 2005; Riedel *et al*., 2011) in sex combs using *42D04-GAL4* and *Laccase2-RNAi* (Dietzl *et al.*, 2007). Laccase2 is required to oxidize dopamine into dopamine quinones and thus acts upstream of Yellow in the melanin synthesis pathway (Figure 4A; Riedel *et al*., 2011). Males with *Laccase2* suppressed in sex combs lacked both black and brown dopamine melanin, making these sex combs appear translucent (Figure 4B). These males displayed strongly reduced mating success compared with wild-type males (Figure 4C) and behavioral defects similar to those observed for *y*^1^ mutants (Movies 9,10), including inefficient grasping of the female for mounting and copulation. We noticed, however, that flies with *Laccase2-RNAi* driven by *42D04-GAL4* also showed a loss of melanin in the aedeagus (Supplementary Figure S6A), which is the main part of the male genitalia used for copulation, despite no visible expression of *42D04-GAL4* in the adult male genitalia (Supplementary Figure S4G) nor changes in aedeagus pigmentation in *y*^1^ mutants (Supplementary Figure 6A). We therefore used subsets of the *42D04* enhancer (Supplementary Figure S6B) to drive expression of *Laccase2-RNAi*, separating the effects of expression in the sex combs from expression in the genitalia (Supplementary Figure S6C). Male mating success was reduced when *Laccase2* suppression reduced melanization in the sex combs, but not the genitalia (Supplementary Figure S6D-G).

**Fig. 4.**
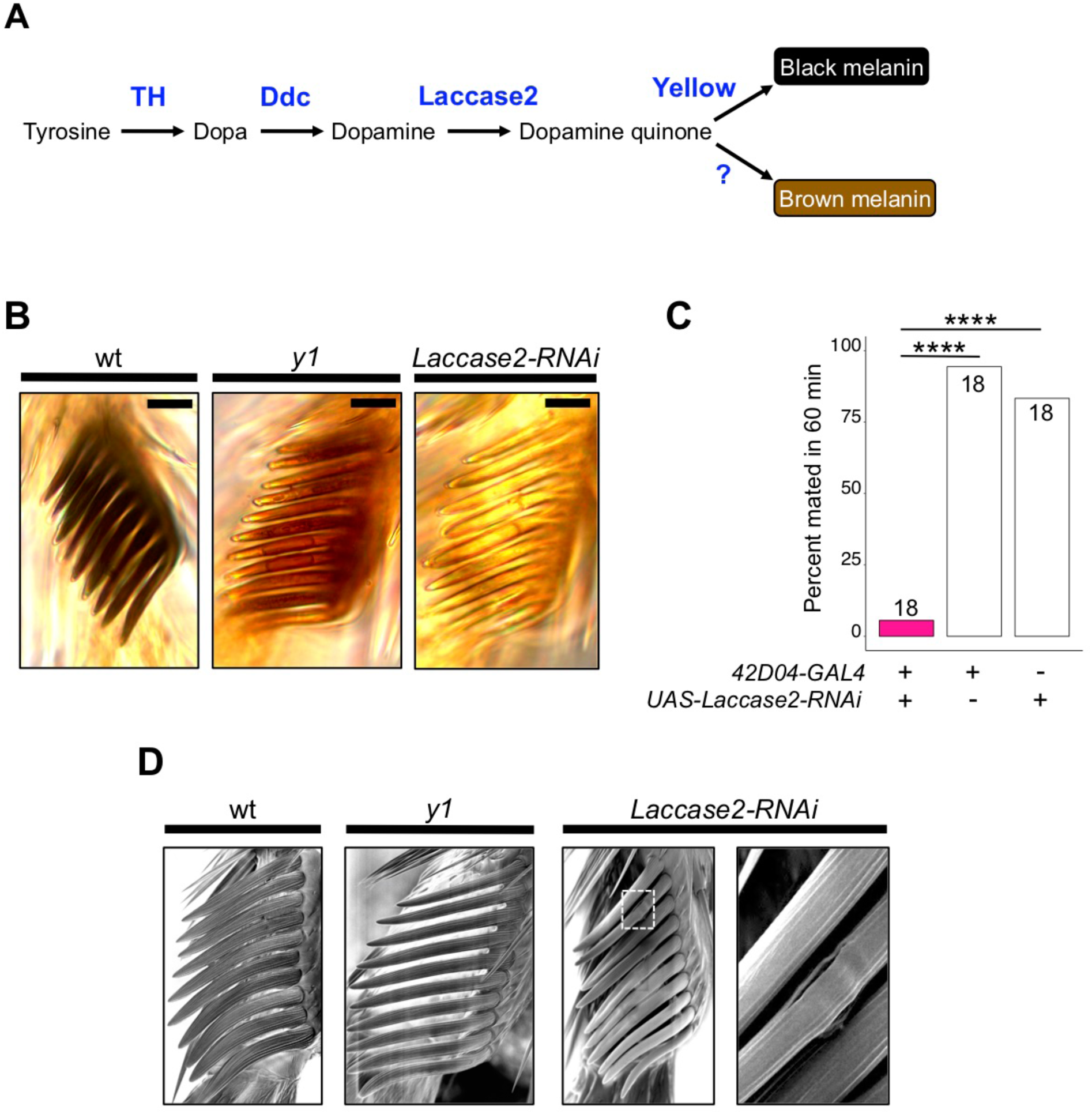
Sex comb melanization is specifically required for male mating success. (**A**) Simplified version of the insect melanin synthesis pathway. (**B**) Light microscopy images of sex combs from wild-type (wt), *y1*, and *42D04-GAL4; UAS-Laccase2-RNAi* males. Expressing *Laccase2-RNAi* in sex combs completely blocked melanin synthesis. (**C**) Expressing *Laccase2-RNAi* using *42D04-GAL4* in males significantly inhibited male mating success. (**D**) Scanning Electron Microscopy (SEM) of sex coms from wild-type (wt), *y1*, and *Laccase2-RNAi* males (expressed using *42D04-GAL4*). Compared to wild-type, sex comb teeth in *y1* mutants appeared thinner and smoother, whereas*Laccase2-RNAi* sex comb teeth appeared even smoother than *y1* mutants, and one comb tooth had a visible crack in the cuticle (white rectangle, enlarged on the right). Scale bars in (**B**) measure 12.5 μm. Sample sizes are shown at the top of each barplot. Significance in was measured using Chi-square tests with Bonferroni corrections for multiple comparisons. \*\*\*\**P*<0.0001.

How can sex comb melanization affect sex comb function? In insects, melanization impacts not only the color of the adult cuticle but also its mechanical stiffness (Xu et al., 1997; Kerwin et al., 1999; Vincent and Wegst, 2004; Anderson 2005; Arakane *et al*., 2005; Suderman *et al*., 2006; Riedel *et al*., 2011; Noh *et al*., 2016). For example, expressing *Laccase2-RNAi* in *D. melanogaster* wings softens the cuticle to such a degree that the wings collapse (Riedel *et al*., 2011). Butterflies lacking dopamine melanin due to loss of *yellow* or another gene required for melanin synthesis, *Dopa decarboxylase*, show changes in the fine structure of their wing scales (Matsuoka and Monteiro, 2018), and we also observed structural changes in *D. melanogaster* sex comb teeth lacking *yellow* or *Laccase2* expression using scanning electron microscopy (SEM), with a crack appearing in one of the *Laccase2-RNAi* comb teeth (Figure 4D). We conclude that these structural changes in sex combs are responsible for inhibiting the *yellow* mutant male’s ability to grasp a female for mounting and copulation (Movie 10). Interestingly, Wilson *et al* (1976) also proposed “that there may be a structural basis for the behavioural effects of the [*yellow*] mutant” based on their observations of behavior in *yellow* mutant males.

Data from other Drosophila species are also consistent with this structural hypothesis. Specifically, *yellow* mutants in *D. subobscura, D. pseudoobscura*, and *D. gaucha*, all of which have sex combs, show reduced male mating success (Rendel, 1944; Tan, 1946; Frias and Lamborot, 1970; Pruzan-Hotchkiss *et al*., 1992) whereas *yellow* mutants in *Drosophila willistoni*, a species that lacks sex combs (Kopp, 2011; Atallah *et al*., 2014), do not (Da Silva *et al*., 2005). Sex comb morphology is highly diverse among species that have sex combs (Kopp, 2011), but these structures generally seem to be melanized (Supplementary Figure S7; Tanaka *et al*., 2009) and used to grasp females (Movies 11-15). Our high-speed video recordings of mating in *D. anannasae, D. bipectinata, D. kikkawai, D. malerkotiana*, and *D. takahashi* show that differences in sex comb morphology (Supplementary Figure S7) correspond with differences in how (where on the female and with which part of the male leg) the male grasps the female prior to copulation (Movies 11-15). It remains unclear how *D. willistoni* males (and males of other species without sex combs) are able to efficiently grasp females prior to copulation (Movie 16).

### Conclusion

Taken together, our data show that melanization of a secondary sexual structure affects mating in *D. melanogaster*. Specifically, we find that the reduced mating success of *D. melanogaster yellow* mutant males, which was perceived as a behavioral defect for decades, is caused by changes in the morphology of the structures used during mating. These observations underscore that behavior cannot be understood by studying the nervous system alone; anatomy and behavior function and evolve as an interconnected system.

## Materials and Methods

### Fly stocks and maintenance

The following lines were used for this work: *y1* [which was backcrossed into a wild-type (*Canton-S*) line for 6 generations before starting our experiments; the *y1* allele contains an A to C transversion in the ATG initiation and is considered a null allele (Geyer *et al*., 1990)]; *Canton-S* as wild-type (courtesy of Scott Pletcher); *UAS-yellow-RNAi* obtained from the Vienna Drosophila Resource Centre (VDRC) (Dietzl *et al.*, 2007, KK106068); *y1;UAS-y* (BDSC 3043); *elav-GAL4* (BDSC 49226); *nsyb-GAL4* (BDSC 39171); *repo-GAL4* (BDSC 7415); *dsx*^*GAL4*^ (Robinett *et al*., 2010) (courtesy of Bruce Baker); *dsx*^*GAL4*^ (Rideout *et al*., 2010) (courtesy of Stephen Goodwin); *fru*^*GAL4*^ (Stockinger *et al*., 2005) (courtesy of Barry Dickson); the following Janelia enhancer trap GAL4 lines (Pfeiffer *et al*., 2008): *40A05-GAL4* (BDSC 48138), *41D01-GAL4* (BDSC 50123), *42D02-GAL4* (BDSC 41250), *41F06-GAL4* (BDSC 47584), *41A01-GAL4* (BDSC 39425), *42D04-GAL4* (BDSC 47588), *40F03-GAL4* (BDSC 47355), *39E06-GAL4* (BDSC 50051), *42C06-GAL4* (BDSC 50150), *40F04* (BDSC 50094); *y*^*mCherry*^ (courtesy of Nicolas Gompel); *nsyb-GAL80* (courtesy of Julie Simpson); *UAS-Laccase2-RNAi* obtained from the VDRC (Dietzl *et al.*, 2007, KK101687); *dsx*^*GAL4-DBD*^ (Pavlou *et al*., 2016) (courtesy of Stephen Goodwin); *vGlut*^*dVP16-AD*^ (Gao *et al*., 2008) (courtesy of Stephen Goodwin); BDSC 6993; BDSC 49365; BDSC 6927; BDSC 45175; BDSC 3740; BDSC 5820; BDSC 8848 (courtesy of Shinya Yamamoto); BDSC 7010 (courtesy of Shinya Yamamoto); *TPH-GAL4* (courtesy of Shinya Yamamoto); *wing-body-GAL4* (BDSC 44373); *D. melanogaster yellow 5’ up EGFP reporter* (Kalay and Wittkopp, 2010) (courtesy of Gizem Kalay); *D. melanogaster yellow intron EGFP reporter* (Kalay and Wittkopp, 2010) (courtesy of Gizem Kalay); *vasa-Cas9* (BDSC 51324); *UAS-cytGFP* (courtesy of Janelia Fly Core); *pJFRC12-10XUAS-IVS-myr::GFP* (courtesy of Janelia Fly Core). All flies were grown at 23°C with a 12 h light-dark cycle with lights on at 8AM and off at 8PM on standard corn-meal fly medium.

### Behavior

#### Mating assays

Virgin males and females were separated upon eclosion and aged for 4-7 d before each experiment. Experiments were carried out at 23°C on a 12 h light dark cycle with lights on at 8 AM and off at 8 PM on standard corn-meal fly medium. Males were isolated in glass vials, and females were group housed in standard plastic fly vials at densities of 20-30 flies. All mating assays were performed at 23°C between 8-11AM or 6-9PM. For each assay replicate, a single virgin male and female fly were gently aspirated into a 35 mm diameter Petri dish (Genesee Scientific, catalog #32-103) placed on top of a 17 inch LED light pad (HUION L4S) and immediately monitored for 60 min for courtship and copulation activity. All genotypes tested initiated courtship (including tapping, chasing, wing extension, genital licking, and attempted copulation) towards the female. Any genotype that copulated within the 60 min window was noted. Except for the experiment described in Figure 5, all female targets in mating assays were wild-type (*Canton-S*). Percent mated in 60 min was then calculated as the number of replicates that mated divided by the total number of replicates and multiplied by 100.

#### Courtship analysis

For courtship analysis, 60 min videos were recorded using Canon VIXIA HF R500 camcorders mounted to Manfrotto (MKCOMPACTACN-BK) aluminum tripods. To calculate courtship indices in Figure 1 between wild-type and *y1* males, the amount of time males spent engaged in courtship: tapping, chasing, wing extension, genital licking, or attempted copulation was quantified for the first 10 min of the assay and divided by the total 10 min period. We chose to quantify courtship activity within the first 10 min of the assay, because wild-type (*Canton-S*) males will often begin copulating after this window, while *y1* males will continue to court throughout the entire 60 min period. Wing extension bouts were quantified by noting every unilateral wing extension bout for each genotype within the first 10 min of the assay.

#### Song analysis

Courtship song was recorded as described previously (Arthur *et* al., 2013). All genotypes were recorded simultaneously. Song data was segmented (Arthur *et al*., 2013) and analyzed (http://www.github.com/dstern/BatchSongAnalysis) without human intervention. *P*-values for one-way ANOVAs were estimated with 10,000 permutations (http://www.mathworks.com/matlabcentral/fileexchange/44307-randanova1).

#### High-speed video capture

For high-speed video capture of attempted mounting and copulation events, virgin males and females were isolated upon eclosion and aged for 4-7 d before each assay. Using a Fascam Photron SA4 (courtesy of Gwyneth Card) mounted with a 105 mm AF Micro Nikkor Nikon lens (courtesy of Gwyneth Card), we recorded individual pairs of males and females that were gently aspirated into a single well of a 96 well cell culture plate (Corning 05-539-200) partially filled with 2% agarose and covered with a glass coverslip. We recorded mounting and copulation attempts at 1000 frames per second (fps) and played back at 30 fps. Most wild-type males attempted mounting 3-5 times before copulating, whereas *y1, yellow-RNAi*, and *Laccasse2-RNAi* males repeatedly attempted mounting without engaging in copulation, mirroring the videos we captured on the Canon VIXIA HF R500 at 30 fps.

### Imaging sex combs and genitalia

Sex comb images highlighting different melanization states (Figure 3I, J, O; Figure 4B) were taken using a Zeiss Axio Cam ERc 5s mounted on a Zeiss Axio Observer A1 Inverted Microscope. Front legs were cut and placed sex comb side down on a microscope slide (Fisher brand 12-550-123) and imaged through a 40x objective. Images were processed using AxioVision LE software. Abdomens and genitalia images highlighting different melanization states of the aedeagus and female genital bristles were captured using a Canon EOS Rebel T6 camera mounted with a Canon MP-E 65 mm macro lens. Genitalia images were processed in Adobe Photoshop (version 19.1.5) (Adobe Systems Inc., San Jose, CA).

Focus Ion Beam Scanning Electron Microscope (FIB-SEM) images (Figure 4D) were taken by placing individual, dissected legs on carbon tape adhered to a SEM pin stud mount with sex combs facing up. The samples were then coated with a 20-nm Au layer using a Gatan 682 Precision Etching and Coating System, and imaged by SEM in a Zeiss Sigma system. The samples were imaged using a 3-nA electron beam with 1.5 kV landing energy at 2.5MHz.

### Immunohistochemistry and confocal imaging

#### Central Nervous System

Dissections, immunohistochemistry, and imaging of fly central nervous systems were done as previously described (Aso *et al.*, 2014). In brief, brains and VNCs were dissected in Schneider’s insect medium and fixed in 2% paraformaldehyde (diluted in the same medium) at room temperature for 55 min. Tissues were washed in PBT (0.5% Triton X-100 in phosphate buffered saline) and blocked using 5% normal goat serum before incubation with antibodies. Tissues expressing GFP were stained with rabbit anti-GFP (ThermoFisher Scientific A-11122, 1:1000) and mouse anti-BRP hybridoma supernatant (nc82, Developmental Studies Hybridoma Bank, Univ. Iowa, 1:30), followed by Alexa Fluor® 488-conjugated goat anti-rabbit and Alexa Fluor® 568-conjugated goat anti-mouse antibodies (ThermoFisher Scientific A-11034 and A-11031), respectively. Tissues expressing mCherry-tagged Yellow protein (*y*^*mCherry*^) were stained with rabbit anti-dsRed (Clontech 632496, 1:1000) and rat anti-*D*N-Cadherin (DN-Ex #8, Developmental Studies Hybridoma Bank, Univ. Iowa, 1:100) as neuropil marker, followed by Cy^TM^3-conjugated goat anti-rabbit and Cy^TM^5-conjugated goat anti-rat antibodies (Jackson ImmunoResearch 111-165-144 and 112-175-167), respectively. After staining and post-fixation in 4% paraformaldehyde, tissues were mounted on poly-L-lysine-coated cover slips, cleared, and embedded in DPX as described. Image z-stacks were collected at 1 μm intervals using an LSM710 confocal microscope (Zeiss, Germany) fitted with a Plan-Apochromat 20x/ 0.8 M27 objective. Images were processed in Fiji (http://fiji.sc/) and Adobe Photoshop (version 19.1.5) (Adobe Systems Inc., San Jose, CA).

#### Sex combs and genitalia

Adult flies were 2-7 d old and pupae were 96 h old after pupal formation (APF) for the EGFP reporter experiment summarized in Supplementary Figure S11. Flies were anesthetized on ice, submerged in 70% ethanol, rinsed twice in phosphate buffered saline with 0.1 % Triton X-100 (PBS-T), and fixed in 2% formaldehyde in PBS-T. Forelegs and genitalia/abdomen tips were removed with fine scissors and mounted in Tris-buffered (pH 8.0) 80% glycerol. Serial optical sections were obtained at 1.5 µm or 0.5 µm intervals on a Zeiss 880 confocal microscope with a LD-LCI 25x/0.8 NA objective (genitalia) or a Plan-Apochromat 40x/1.3 NA objective (appendages/tarsal sex combs). The native fluorescence of GFP, mCherry and autofluorescence of cuticle were imaged using 488, 594 and 633 lasers, respectively. Images were processed in Fiji (http://fiji.sc/), Icy (http://icy.bioimageanalysis.org/) and Adobe Photoshop (version 19.1.5) (Adobe Systems Inc., San Jose, CA).

### Statistics

Statistical tests were performed in R for Mac version 3.3.3 (R Core Team 2018) using Fisher’s exact tests to test for statistically significant effects of 2 × 2 contingency tables, Chi-square tests to test for statistically significant effects of contingency tables greater than 2 × 2 with Bonferroni corrections for multiple comparisons, and two-tailed Student’s t-tests to test for statistically significant effects of pairwise comparisons of continuous data with normally distributed error terms. For song analysis, one-way ANOVAs were performed in MATLAB version R2017a (The MathWorks, Inc.).

### Generation of the mating regulatory sequence (MRS) deletion line

Using the 209 bp region mapped in Drapeau *et al*. (2006) between −300 and −91 bp upstream of *yellow*’s transcription start site, we designed two single guide RNA (gRNA) target sites at −291 bp and −140 bp that maximized the MRS deletion region, given constraints of identifying NGG PAM sites required for CRISPR/Cas9 gene editing (Supplementary Figure S2F). We in-vitro transcribed these gRNAs using a MEGAscript T7 Transcription Kit (Invitrogen) following the PCR-based protocol from Bassett *et al*. (2013). Two 1 kb homology arms were PCR amplified from the *yellow* locus immediately upstream and downstream of the gRNA target sites using the forward and reverse primers with NcoI and BglII tails, respectively, for the Left Arm (5’-TTACCATGGGGGATCAAGTTGAACCAC-3’, 5’-GGAGATCTGGCCTTCATCGACATTTA-3’) and the forward and reverse primers with Bsu36I and MluI tails, respectively, for the Right Arm (5’-TACATCCCTAAGGCCTGATTACCCGAACACT-3’, 5’-TATACGCGTTGCCATGCTATTGGCTTC-3’) and cloned into pHD-DsRed-attp (Gratz *et al*., 2014; Addgene Plasmid # 51019) in two steps, digesting first with NcoI and BglII (Left Arm) to transform the Left Arm and second with Bsu36I and MluI (Right Arm) to transform the Right Arm, flanking the 3xP3::DsRed, attP, and LoxP sites. Homology arms were ligated into pHD-DsRed-attp using T4 DNA Ligase (ThermoFisher Scientific), and products were transformed into One Shot TOP10 (Invitrogen) DH5 alpha competent cells. Purified donor plasmid was then co-injected at 500 ng/uL with the two gRNAs at 100 ng/uL total concentration into a *vasa-Cas9* (BDSC 51324) line. Flies were then screened for DsRed expression in the eyes, and Sanger sequenced verified for a 3xP3::DsRed replacement of the MRS region (Supplementary Figure S2F). We confirmed that we deleted 152 bp of the 209 bp region based on Sanger sequencing the CRISPR/Cas9 cut sites (Supplementary Figure S2F). Next, we crossed *y*^*δMRS+3xP3::DsRed*^ with a Cre-expressing fly line (courtesy of Bing Ye, University of Michigan) to excise 3xP3::DsRed and screened for flies that lost DsRed expression in the eyes. Finally, we PCR-gel verified that DsRed was indeed removed in creation of the *y*^*δMRS*^ line using the forward and reverse primers, respectively (5’-CAGTCGCCGATAAAGATGAACACTG-3’, 5’-CAAGGTGATCAGGGTCACAAGGATC-3’) (Supplementary Figure S2G).

### Generation of the 42D04-GAL4 enhancer sub-fragment pBPGUw lines

Enhancer sub-fragments (2 kb, 2 kb, 1.3 kb, 1.3 kb, and 1.3 kb for *42D04_A,B,C,D,E*-GAL4, respectively) were synthesized as IDT gene blocks (sequences available in Supplementary File S1) based off of the 42D04 *D. melanogaster dsx* enhancer sequence (FBsf0000164494) (Supplementary Figure S7). The gene blocks were designed with 5’ and 3’ Gibson tails to facilitate Gibson assembly (Gibson *et al*., 2009) into the GAL4 plasmid pBPGUw (Pfeiffer *et al*., 2008; Addgene Plasmid #17575) after digestion with FseI and AatII. Products were transformed into Mix and Go! DH5 alpha competent cells (Zymo). Clones were selected by ampicillin resistance on Amp-LB plates (60mg/mL). Purified plasmids were injected at 500 ng/uL into the phiC31 integrase-expressing 86Fb landing site line *BDSC 24749* (courtesy of Rainbow Transgenics) for phiC31 attP-attB integration and screened for using a mini-white marker.

## Supporting information

Movie_S1_Canton_S_short_2

Movie_S2_y1_short

Movie_S3_wt_copulation_success

Movie_S4_y1_extended_copulation_fail

Movie_S5_dsx_Goodwin_yellow-RNAi

Movie_S6_42D04>yellow-RNAi

Movie_S7_y1_1000fps_1

Movie_S8_wt_1000fps

Movie_S9_42D04>laccase2-RNAi

Movie_S10_42D04>laccase2-RNAi_1000fps

Movie_S11_D_anannasae

Movie_S12_D_bipectinata

Movie_S13_D_kikkawai

Movie_S14_D_malerkotiana

Movie_S15_D_takahashi

Movie_S16_D_willistoni

## Acknowledgments

We thank members of the Wittkopp and Stern labs for helpful discussions. For fly strains, we thank Bruce Baker, Carmen Robinett, Stephen Goodwin, Barry Dickson, Scott Pletcher, Julie Simpson, Shinya Yamamoto, Bing Ye, Nicolas Gompel, Gizem Kalay, The Bloomington Drosophila Stock Center, The Vienna Drosophila RNAi Center, and the Janelia Fly Core for fly strains. For fly injections, we thank Rainbow Transgenics Inc. For technical support with Scanning Electron Microscopy (SEM), we thank Harald Hess and Song Pang. For use of the Photron for high-speed video capture, we thank Gwyneth Card and W. Ryan Williamson. CNS dissections, immunostaining, and imaging were performed by the Janelia Project Technical Resource team with special thanks to Gudrun Ihrke, Kari Close, and Christina Christoforou. We thank Nicolas Gompel, Abby Lamb, and Henry Ertl for comments on the manuscript.

## Data and materials availability

All data is available in the main text or the supplementary materials.

## Supplementary Figures

**Supplemental Figure S1.**
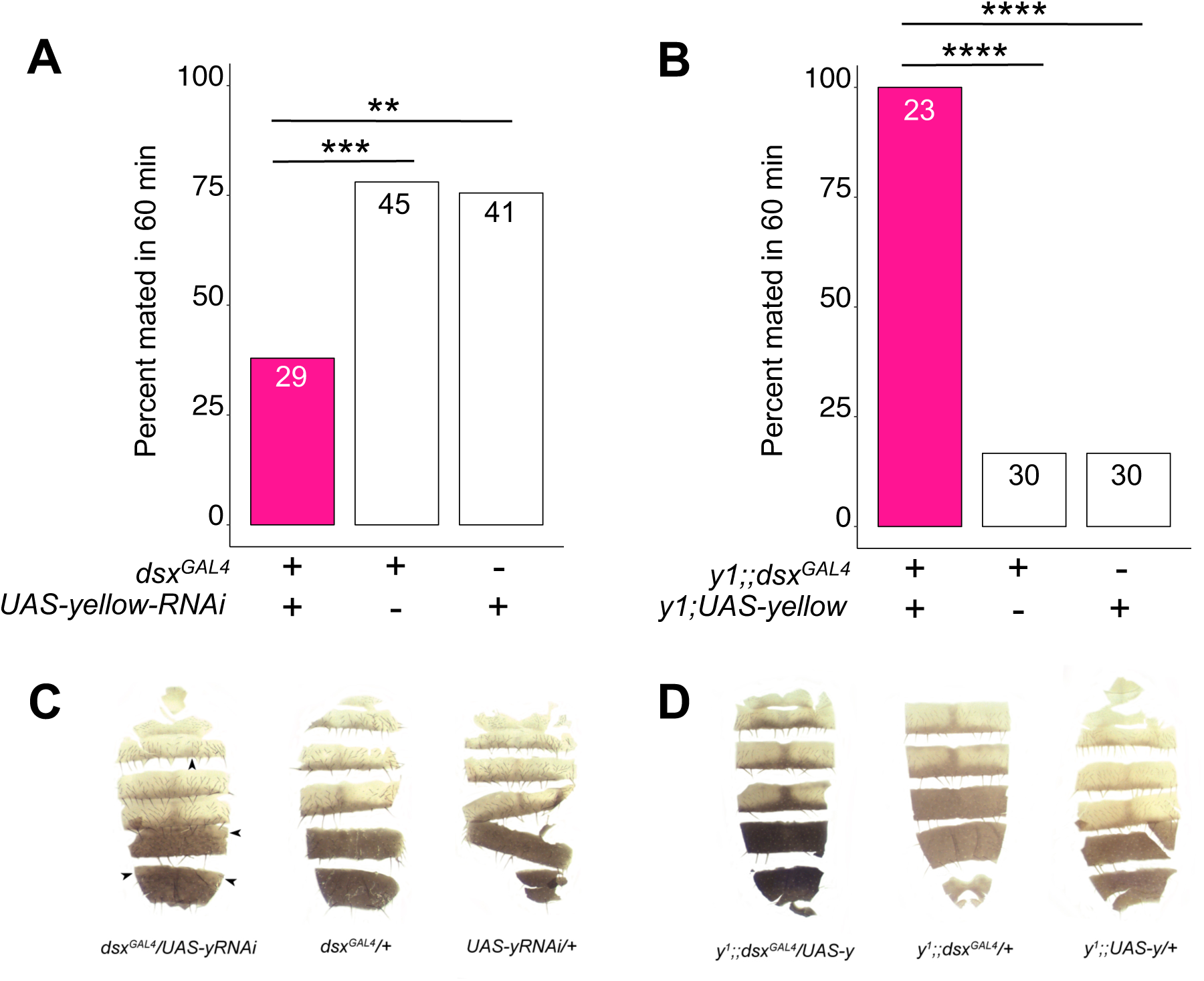
*yellow* expression in *dsx*-expressing cells is necessary and sufficient for male mating success. (**A**) Expressing *yellow-RNAi* in *dsx*-expressing cells using *dsx*^*GAL4*^ (Rideout *et al*., 2010) significantly inhibited male mating success. (**B**) Expressing *yellow* in *dsx*-expressing cells using *dsx*^*GAL4*^ in a *y1* mutant background was sufficient to restore male mating success. (**C**) Expressing *yellow-RNAi* using *dsx*^*GAL4*^ (Rideout *et al.*, 2010) partially reduced black melanin levels in the male A5 and A6 abdominal tergites, consistent with prior work (Williams *et al*. 2008, Rogers *et al*. 2014, Kalay *et al*. 2016). (**D**) Expressing *yellow* using *dsx*^*GAL4*^ partially elevated black melanin levels in the male A5 and A6 abdominal tergites. Sample sizes are shown at the top of each barplot. Significance was measured using Chi-square tests with Bonferroni corrections for multiple comparisons. \*\**P*<0.01, \*\*\**P*<0.001, \*\*\*\**P*<0.001.

**Supplemental Figure S2.**
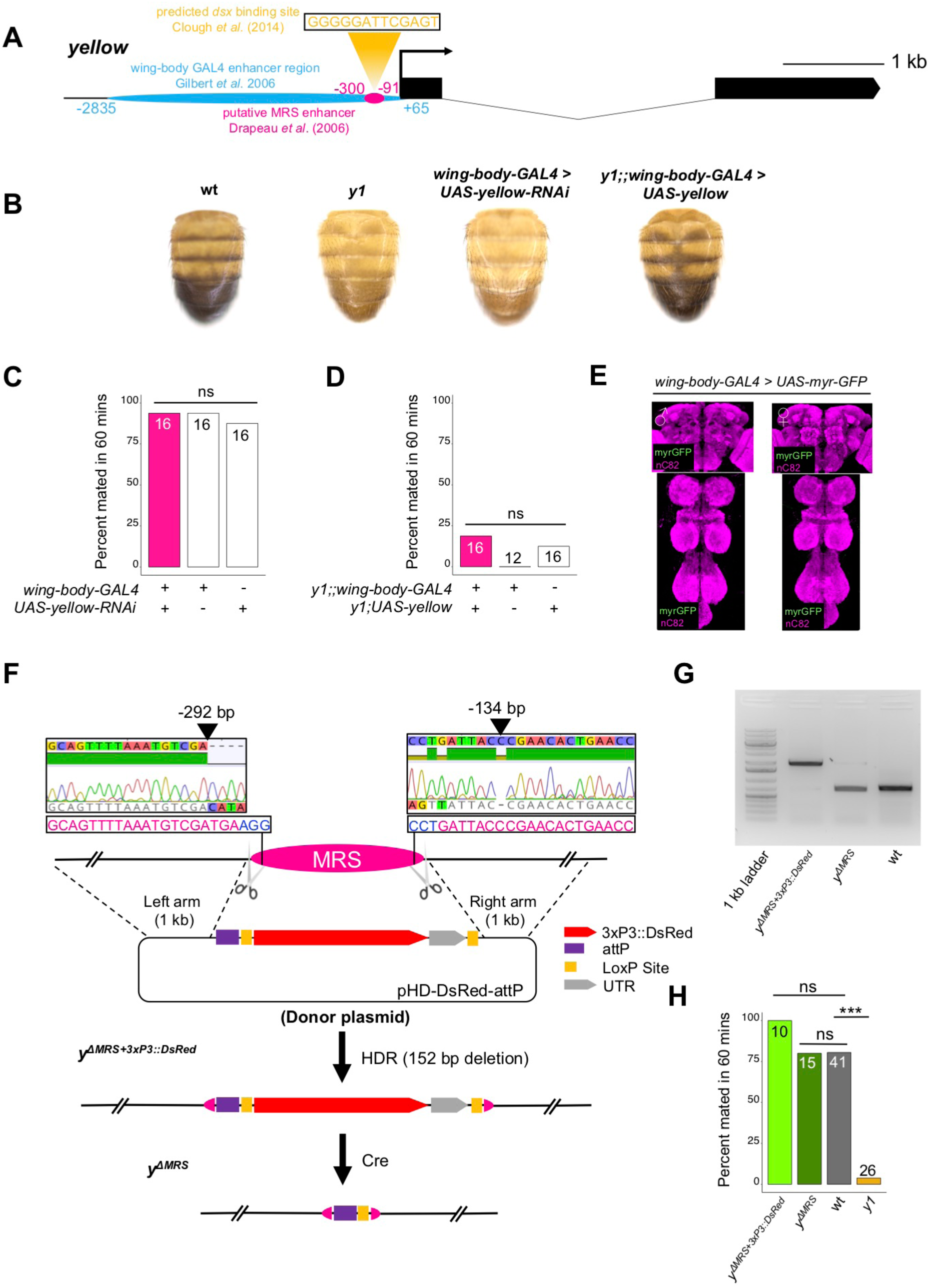
The mating regulatory sequence (MRS) from Drapeau *et al*. (2006) does not affect male mating success. (**A**) Diagram of the *yellow* locus highlighting the putative “mating regulatory sequence” (MRS) (pink) region mapped in Drapeau *et al*. (2006) and a predicted *dsx* binding site (yellow) identified by ChIP-seq in Clough *et al*. (2014). The predicted binding site was identified based on *in vivo* Doublesex occupancy data (PWM score = 88.7) localized between 356,273 and 356,286 bp on the X chromosome (see Supplementary Table S2 in Clough *et al*., 2014). The wing-body enhancer region is indicated in blue, which was cloned upstream of GAL4 in Gilbert *et al*. (2006) to make the *wing-body-GAL4* line. (**B**) Expressing *yellow-*RNAi using *wing-body-GAL4* reduced black melanin to *y1* levels, and expressing *yellow* in a *y1* mutant background using *wing-body-GAL4* restores black melanin synthesis to wild-type (wt) levels. (**C**) Expressing *yellow-*RNAi using *wing-body-GAL4* did not inhibit male mating success. (**D**) Expressing *yellow* using *wing-body-GAL4* in a *y1* mutant background did not restore male mating success. (**E**) Brain and VNC of adult male and female flies stained with anti-GFP (green) antibody for myrGFP expressed using *wing-body-GAL4* and counterstained with anti-nC82 (magenta) for neuropil. (**F**) Diagram illustrating the CRISPR/Cas9-facilitaed homology-directed repair (HDR) strategy used to excise and replace the MRS (pink) with pHD-DsRed-attP (red) (Gratz *et al*., 2014). Two sgRNAs (pink letters) were designed towards target PAM sites (blue letters) at the most 5’ and 3’ bounds of the MRS (scissors). Sanger sequencing chromatograms illustrate the location of each cut site (black arrows) relative to the transcription start site. DsRed was removed using Cre-lox recombinase (Siegal and Hartl 1996). (**G**) PCR validation of DsRed removal and MRS deletion. (**H**) Excising the putative MRS did not inhibit male male mating success. Sample sizes are shown at the top of each barplot. Significance was measured using Chi-square tests with Bonferroni corrections for multiple comparisons. \*\*\**P*<0.001. n.s., not significant.

**Supplemental Figure S3.**
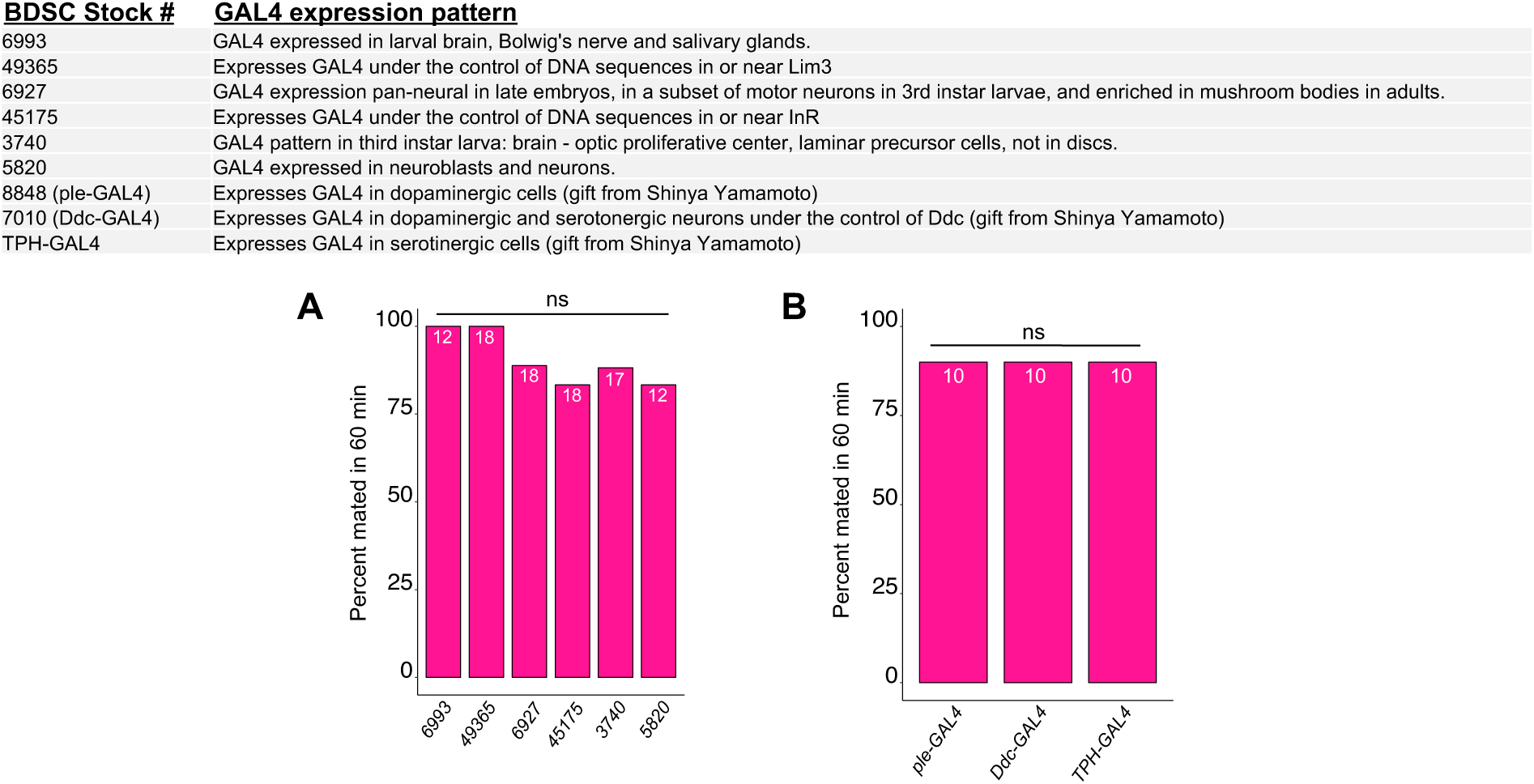
Expressing *yellow-*RNAi in subsets of CNS tissue does not affect male mating success. (**A**,**B**) Expressing *yellow-*RNAi using a series of CNS, dopaminergic, and serotonergic GAL4 drivers did not affect male mating success. Significance was measured using Chi-square tests with Bonferroni corrections for multiple comparisons. Sample sizes are shown at the top of each barplot. Significance was measured using Chi-square tests with Bonferroni corrections for multiple comparisons. n.s., not significant.

**Supplemental Figure S4.**
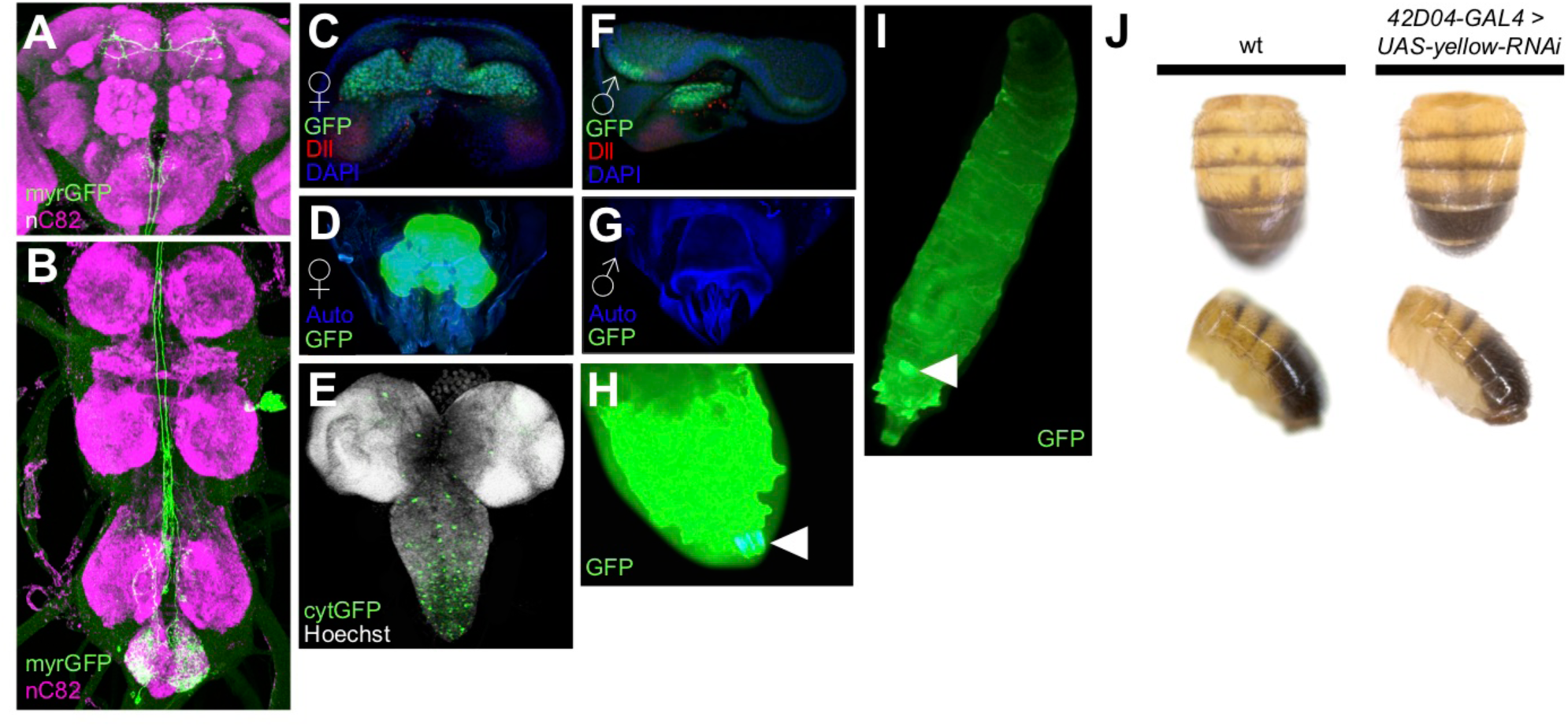
Expression pattern of *42D04-GAL4*. (**A**,**B**) Brain and VNC of adult female fly stained with anti-GFP (green) antibody for myrGFP expressed using *42D04-GAL4* and counterstained with anti-nC82 (magenta) for neuropil. (**C**) L3 larval female genital disc stained with anti-GFP (green) antibody for cytGFP expressed using *42D04-GAL4*, anti-Dll (red) for Distal-less expression, and counterstained with DAPI (blue) for DNA (courtesy of Janelia Fly Light). (**D**) Adult female genitalia native cytGFP (green) expressed using *42D04-GAL4.* (**E**) L3 CNS native cytGFP (green) expressed using *42D04* (**F**) L3 larval male genital disc stained with anti-GFP (green) antibody for cytGFP expressed using *42D04-GAL4*, anti-Dll (red) for Distal-less expression, and counterstained with DAPI (blue) for DNA (courtesty of Janelia Fly Light). (**G**) Adult male genitalia did not show native cytGFP expression using *42D04-GAL4*. (**H**) L3 larval posterior spiracle (white arrowhead) native cytGFP (green) expression. (**I**) L3 larva whole body highlighting native cytGFP (green) expression in the genital disc (white arrowhead). (**J**) Expressing *yellow-*RNAi using *42D04-GAL4* does not affect body pigmentation relative to wild-type (wt) flies.

**Supplemental Figure S5.**
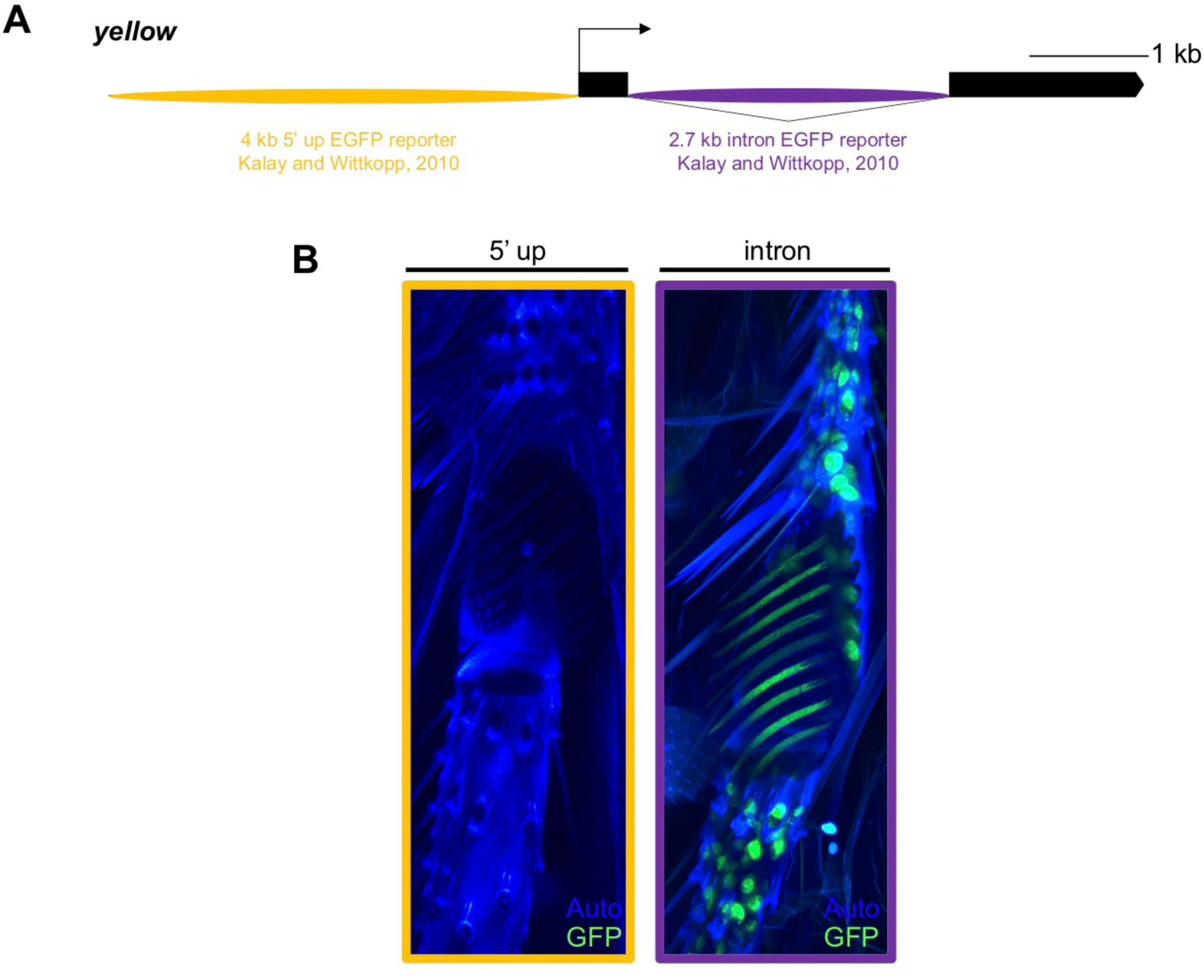
*yellow* EGFP reporters localize *yellow* sex comb expression to the intronic bristle enhancer. (**A**) Diagram of the *yellow* locus highlighting two *D. melanogaster* enhancer regions [5’ up including the wing, body, and putative MRS enhancers reported in Geyer and Corces (1987), Martin *et al*., (1989), and Drapeau *et al*., (2006); and intron, including the bristle and putative sex comb enhancer reported in Geyer and Corces (1987) and Martin *et al*., (1989)] that were cloned upstream of an EGFP reporter in Kalay and Wittkopp (2010). (**B**) Confocal image of a 96 h old (APF) pupal sex comb expressing cytGFP under the control of the 5’ up enhancer region. (**C**) Confocal image of a 96 h APF pupal sex comb expressing cytGFP under the control of the intronic enhancer region, highlighting expression in bristle sockets, sex comb sockets, and sex comb teeth.

**Supplemental Figure S6.**
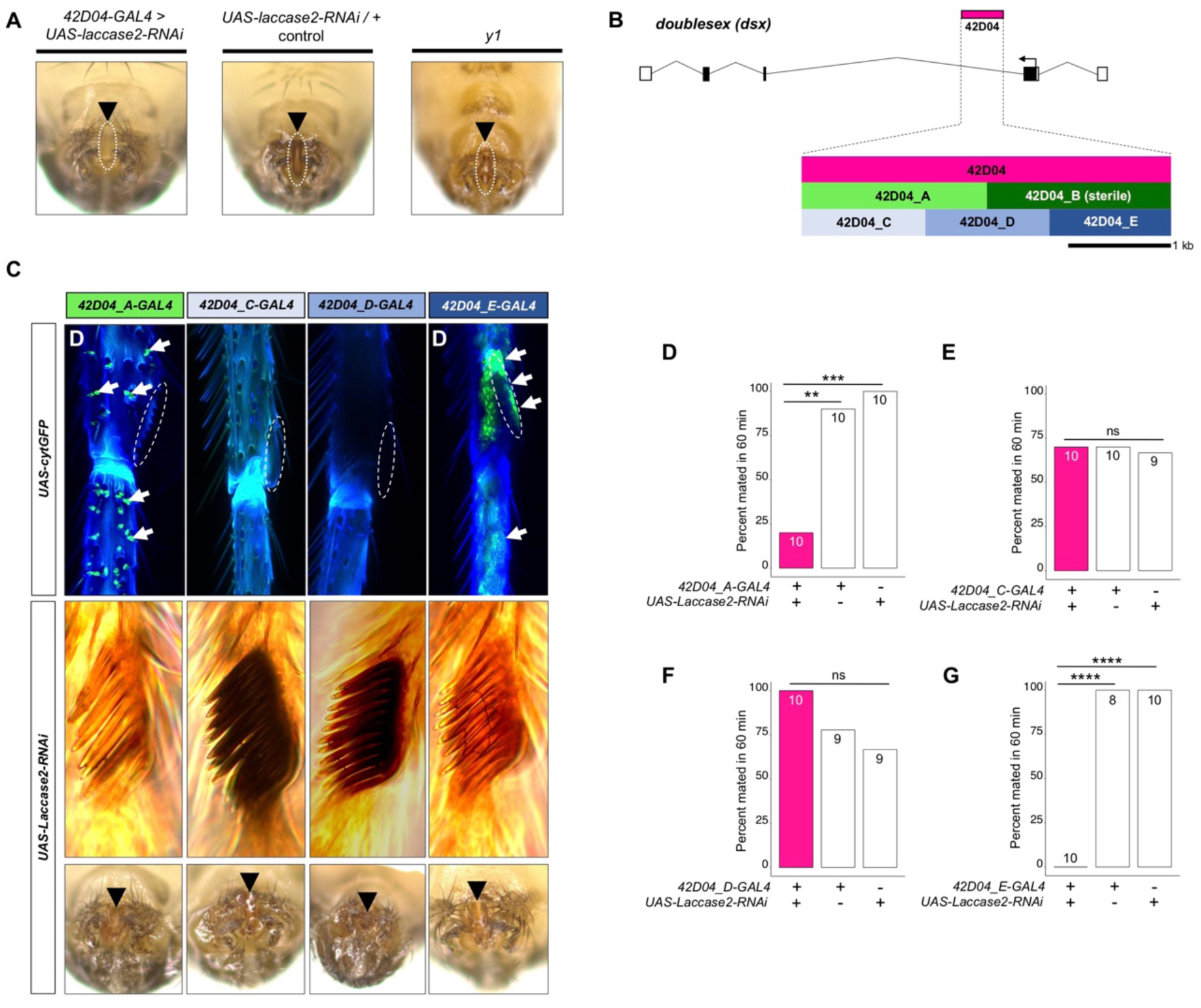
Genetic dissection of the *42D04-GAL4* enhancer confirms the specific role of sex comb melanization, and not the aedeagus, in male mating success. Expressing *Laccase2-RNAi* using *42D04-GAL4* blocked melanin synthesis in the aedeagus. Diagram of the male exon structure of the *dsx* locus highlighting the strategy used to dissect the *42D04-GAL4* expression pattern. Five new GAL4 lines were created by synthesizing different sized sub-fragments of the *42D04-GAL4* enhancer fragment and cloning them upstream of GAL4 (see Supplemental Materials and Methods). Note, *42D04_B-GAL4* could not be maintained, since female flies expressing GAL4 using this enhancer region were all sterile and showed necrotic growths on their genitalia. (**C**) Expression pattern of *42D04_A,C,D, and E-GAL4* lines. Expressing cytGFP using *42D04_A-GAL4* showed GFP (green) localized to bristle sockets, and *42D04_E-GAL4* shows bright GFP in the sex comb and lower leg region. *42D04_C-GAL4* and *42D04_D-GAL4* did not show GFP expression in the legs. Expressing *Laccase2-RNAi* using *42D04_A-GAL4* and *42D04_E-GAL4* blocked melanin synthesis in the sex combs but not the aedeagus. (**D**) Expressing *Laccase2-RNAi* using *42D04_A-GAL4* and *42D04_E-GAL4* inhibited male mating success. Sample sizes are shown at the top of each barplot. Significance was measured using Chi-square tests with Bonferroni corrections for multiple comparisons. \*\**P*<0.01, \*\*\**P*<0.001, \*\*\*\**P*<0.001. n.s., not significant.

**Supplementary Figure S7.**
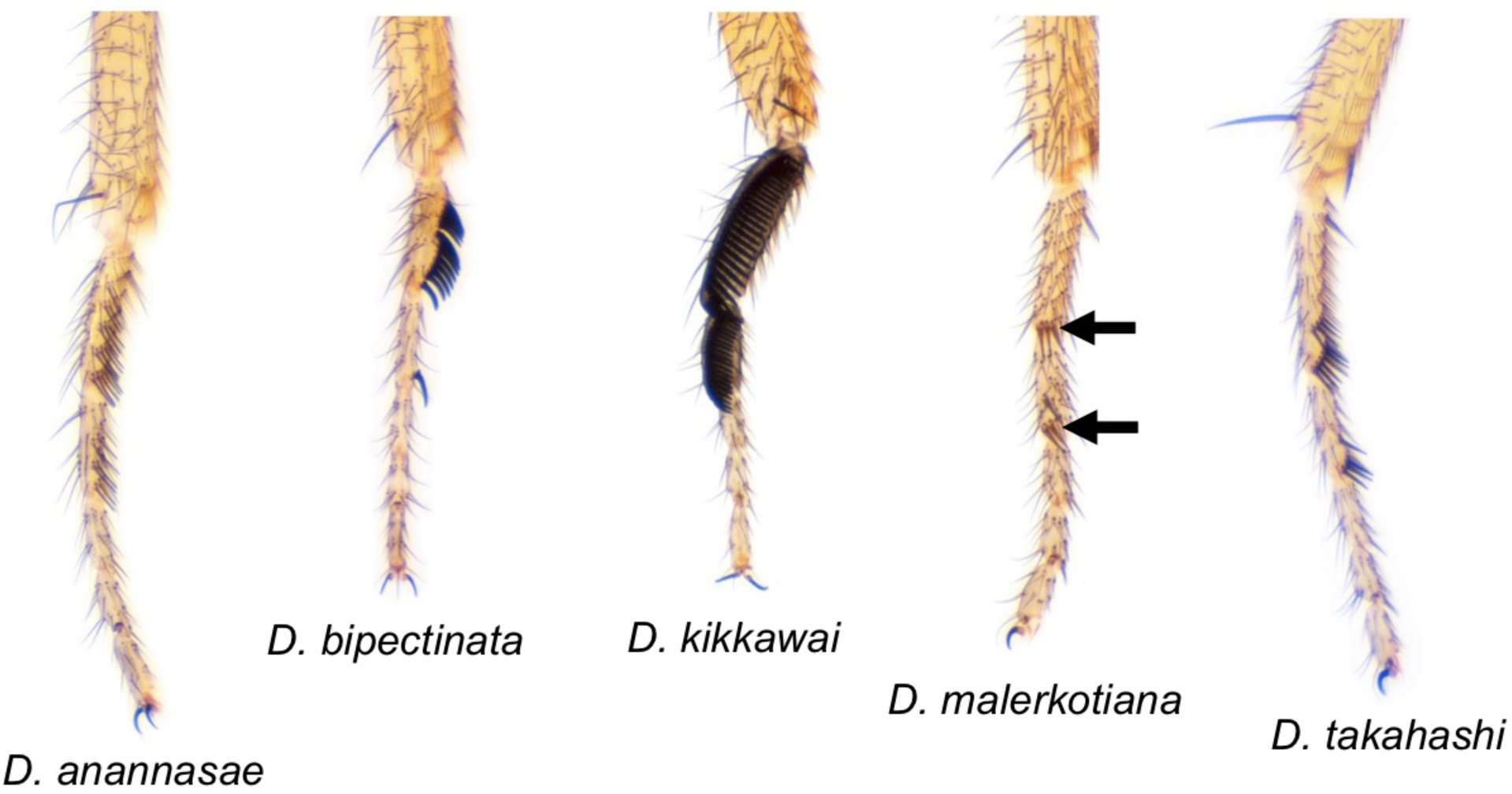
*Drosophila* species with varying sex comb morphology used for high-speed video assays. *anannasae, D. bipectinata, D. kikkawai, D. malerkotiana*, and *D. takahahi* male front forelegs, highlighting variation in sex comb morphology (Nicolas Gompel).

## Movies

**Movie 1. Wild-type courtship and copulation**

**Movie 2. *y1* courtship with wild-type female**

**Movie 3. Wild-type copulation**

**Movie 4. Copulation attempts between *y1* male and wild-type female after 3 h of courtship**

**Movie 5. Copulation attempts between male expressing *yellow-RNAi* in *dsx*^*GAL4*^-expressing cells and wild-type female**

**Movie 6. Copulation attempts between male expressing *yellow-RNAi* in *42D04*-*GAL4*-expressing cells and wild-type female**

**Movie 7. High-speed (1000 fps) video capture of copulation attempts between *y1* male and wild-type female**

**Movie 8. High-speed (1000 fps) video capture of wild-type copulation**

**Movie 9. Copulation attempts between male expressing *Laccase2-RNAi* in *42D04*-*GAL4*-expressing cells and wild-type female**

**Movie 10. High-speed (1000 fps) video capture of copulation attempts between male expressing *Laccase2-RNAi* in *42D04-GAL4*-expressing cells and wild-type female**

**Movie 11. *Drosophila anannasae* wild-type copulation**

**Movie 12. *Drosophila bipectinata* wild-type copulation**

**Movie 13. *Drosophila kikkawai* wild-type copulation**

**Movie 14. *Drosophila malerkotiana* wild-type copulation**

**Movie 15. *Drosophila takahashi* wild-type copulation**

**Movie 16. *Drosophila willistoni* wild-type copulation**

